# SCENITH: A flow cytometry based method for functional profiling energy metabolism with single cell resolution

**DOI:** 10.1101/2020.03.10.985796

**Authors:** Rafael J. Argüello, Alexis J. Combes, Remy Char, Evens Bousiquot, Julien-Paul Gigan, Voahirana Camosseto, Bushra Samad, Jessica Tsui, Peter Yan, Sebastien Boissonneau, Dominique Figarella-Branger, Emeline Tabouret, Evelina Gatti, Matthew F. Krummel, Philippe Pierre

## Abstract

Energetic metabolism reprogramming is critical for cancer and immune responses. Current methods to functionally profile the global metabolic capacities and dependencies of cells are performed in bulk. We designed a simple method for complex metabolic profiling called SCENITH, for Single Cell ENergetIc metabolism by profilIng Translation inHibition. SCENITH allows for the study of metabolic responses in multiple cell types in parallel by flow cytometry. SCENITH is designed to perform metabolic studies ex vivo, particularly for rare cells in whole blood samples, avoiding metabolic biases introduced by culture media. We analyzed myeloid cells in solid tumors from patients and identified variable metabolic profiles, in ways that are not linked to their lineage nor their activation phenotype. SCENITH ability to reveal global metabolic functions and determine complex and linked immune-phenotypes in rare cell subpopulations will contribute to the information needed for evaluating therapeutic responses or patient stratification.

## Introduction

Energetic metabolism (EM) comprises a series of interconnected biochemical pathways capable of using energy rich molecules to produce ATP. In the presence of enough oxygen, cells can produce ATP either by oxidative phosphorylation (OXPHOS) and/or by performing glycolysis. Glycolytic metabolism in the presence of non-limiting concentration of oxygen is called aerobic glycolysis and is characteristic of proliferating cells. Aerobic glycolytic metabolism not only supports proliferation but also cell survival in hypoxic conditions. Immune cells are specially adapted to migrate into peripheral tissues and change of microenvironment. Their energetic metabolism profile is known to correlate with the microanatomical localization, activation, proliferation or functional state (O’Sullivan et al., 2019; Russell et al., 2019). Activation and proliferation drives the differentiation of long lived quiescent naïve into effector T cells. This process is tighly linked to a metabolic switch from OXPHOS to aerobic glycolysis(Pearce et al., 2009; Roos and Loos, 1973). Interestingly, naïve cells re-circulate between the blood and lymph nodes while blood circulating effector cells need to reach peripheral tissues (i.e. infection site or tumors). Supporting the idea that metabolic profile is pre-adapted and is related to cell function, post-mitotic blood immune cells whose function is to rapidly migrate into hypoxic/damaged tissues, such as circulating neutrophils and monocytes, are already engaged in aerobic glycolysis in the blood. Altogether, the current paradigm suggests that a particular metabolic profile of the cells in the blood and tissues is associated to their competence to migrate, proliferate and exert their effector functions in target tissues. However, due to technical limitations and the heterogeneity of cell types, it is not clear which subsets of immune cells rapidly adapt to changing environments or which have predefined metabolic profile.

Competition for glucose within the tumor microenvironment influences cancer progression and the anti-tumoral immune response by regulating EM in both tumoral cells and tumor-infiltrating lymphocytes (TILs), consequently influencing their function(Chang et al., 2015). The presence of effector immune cells into target tissue has clinical relevance and is a well accepted predictive parameter to determine response to immunotherapy(Galon et al., 2006). The success of immunotherapies (e.g. checkpoint inhibitors) is restricted to a relatively small proportion of patients that require a functional and metabolically competent immune system to respond to treament(Antonia et al., 2018; Wolchok et al., 2017). Consequenly, the need of better and accessible tools for patient immuno-metabolic profiling as a strategy for stratification, and monitoring responses to immunotherapies is strongly needed. The metabolic profile of innate and adaptive immune cells correlates with the type of cytokines they produce, and from this point of view metabolic profiles represent a read-out for functional states.

Current methods to determine EM profiles can be classified in three groups. The first group epitomized by Seahorse^®^ (Agilent technologies), uses metabolic inhibitors (i.e. 2-DeoxyGlucose/”DG” and Oligomycin A/”O”) during monitoring of the extracellular acidification rate (ECAR), as well as oxygen consumption rate (OCR) of a large amount of cells in culture, to establish the dependency and metabolic capacities of the cells on different EM pathways. The second group, also a bulk measurement, quantifies the activity of enzymes implicated in particular metabolic pathways in fixed cells or lysates(Miller et al., 2017). Finally, the third group uses mass spectometry and mass-spectrometry imaging to measure the levels of different metabolites produced by EM pathways(Palmer et al., 2016). Very recently, CyTOF-based quantification of different metabolic enzymes was used in parallel to histology to correlate expression of metabolic enzymes with the microanatomical localization of mouse and human T cells(Ahl et al., 2020; Hartmann et al., 2020; Levine et al., 2020).

Current methods need large numbers of highly purified cells(Zhang et al., 2012), and dedicated equipments, and thus cannot be applied to establish functional EM profiles of heterogenous and scarce living cell populations obtained from human samples or biopsies(Llufrio et al., 2018) (Table 1). Given the unpredictability of functional metabolism by phenotyping the levels of a relatively small subset of enzymes in different cell types, we focused on developing a method that functionally quantifies energetic metabolism.

**Table 1.** Comparative table of methods to profile metabolism.

| Method | CyTOF (e.g. Met-Flow, scMEP) | MSI | Seahorse® | SCENITH™ |
| --- | --- | --- | --- | --- |
| Output | Metabolic phenotype | Metabolomic profile (unbiased) | Metabolic capacities and dependencies | Metabolic capacities and dependencies |
| Functional profile of the cells (# of treatments) | NO | NO | YES (4) | YES (non-limited) |
| Cell purification required | NO | NO | YES | NO |
| Single cell resolution | YES | YES | NO | YES |
| Phenotypic analysis | YES | NO | NO | YES |
| Compatible with cell sorting | NO | NO | NO | YES* |
| Ex vivo application | YES | YES | NO | YES |
| Metabolic Readout | Levels of markers (min 10 channels) | Metabolite levels | Changes in extracellular pH and [O<sub>2</sub>] | Changes in protein synthesis levels (one channel) |
| Time (Hs) from sampling to profiling | 0-1 | 0-1 | 24 | 0-1 |
| # cells required in subsets | 500 | 200 | 1,000,000 | 2000 |
| Equipment needed | CyTOF cytometer | Any Imaging Mass cytometer | Seahorse Analyser | Any Flow cytometer# |

\* Not shown

Approximatively half of the total energy that mammalian cells produce by degrading glucose, aminoacids and/or lipids is immediately consumed by the protein synthesis machinery(Buttgereit and Brand, 1995; Lindqvist et al., 2018; Schimmel, 1993). The tremendous energetic cost associated with mRNA translation offers a methodological opportunity of determining the protein synthesis levels as a measure of global EM activity, and its response to environmental cues or different metabolic inhibitors. We took advantage of the drug puromycin, whose incorporation is a highly reliable readout for measuring protein synthesis levels(Andrews and Tata, 1971; Miyamoto-Sato et al., 2000; Nemoto et al., 1999; Wool and Kurihara, 1967), combined to a novel anti-puromycin monoclonal antibody, to develop a reliable method to perform EM profiling with single cell level resolution based on protein synthesis intensity as read-out. We termed this method SCENITH, with reference to our previous SUnSET(Schmidt et al., 2009) and SunRiSE(Argüello et al., 2018) methods for studying translation. SCENITH was used to deconvolve the complex energetics of blood T cells and human tumors-associated myeloid cells, with a much-needed resolution for analysing these highly heterogenous samples, from which the circuitry for metabolism, particularly amongst immune cell subsets, has remained inaccessible.

## Results

### Characterizing the energetic metabolism profile by monitoring changes in protein synthesis levels in response to metabolic inhibitors

To test whether the level of protein synthesis (PS) and the pool of ATP are tightly kinetically coupled, we measured in mouse embryonic fibroblasts (MEF), on the one hand ATP levels using the CellTiter-Glo^®^ luminescence based system, and on the other hand PS levels via puromycin incorporation and staining, after blocking ATP production (Fig. 1a). To completely inhibit ATP production, we treated cells with a mix of inhibitors that block both glycolysis and oxidative phosphorylation (OXPHOS); 100mM 2-Deoxy-D-Glucose (DG), 1μM FCCP and 1μM Oligomycin A (O) (Fig. 1a). To further optimize the signal to noise ratio of puromycin intracellular detection, we developed a novel monoclonal anti-puromycin antibody (clone R4743L-E8, Rat IgG2a) specifically adapted to intracellular flow cytometry. Both protein synthesis levels (Fig. 1b,d) and ATP levels (Fig. 1c) dropped within 5-10 minutes after blocking ATP synthesis, with a strikingly similar slope, showing that changes in ATP levels and PS levels are tighly correlated (Fig. 1e; r 0.985; P<0.0001). To test the relationship between ATP consumption and translational and transcriptional activities, we treated cells with the same inhibitors to block de novo ATP synthesis, together with translation and/or transcription inhibitors. Altogether, our results confirmed that protein synthesis is one of the most energy consuming metabolic activities (Supplementary fig. 1), and most importantly, it represents a stable and reliable readout to evaluate rapidly the impact of metabolic pathways inhibition on the cell.

**Figure 1.**
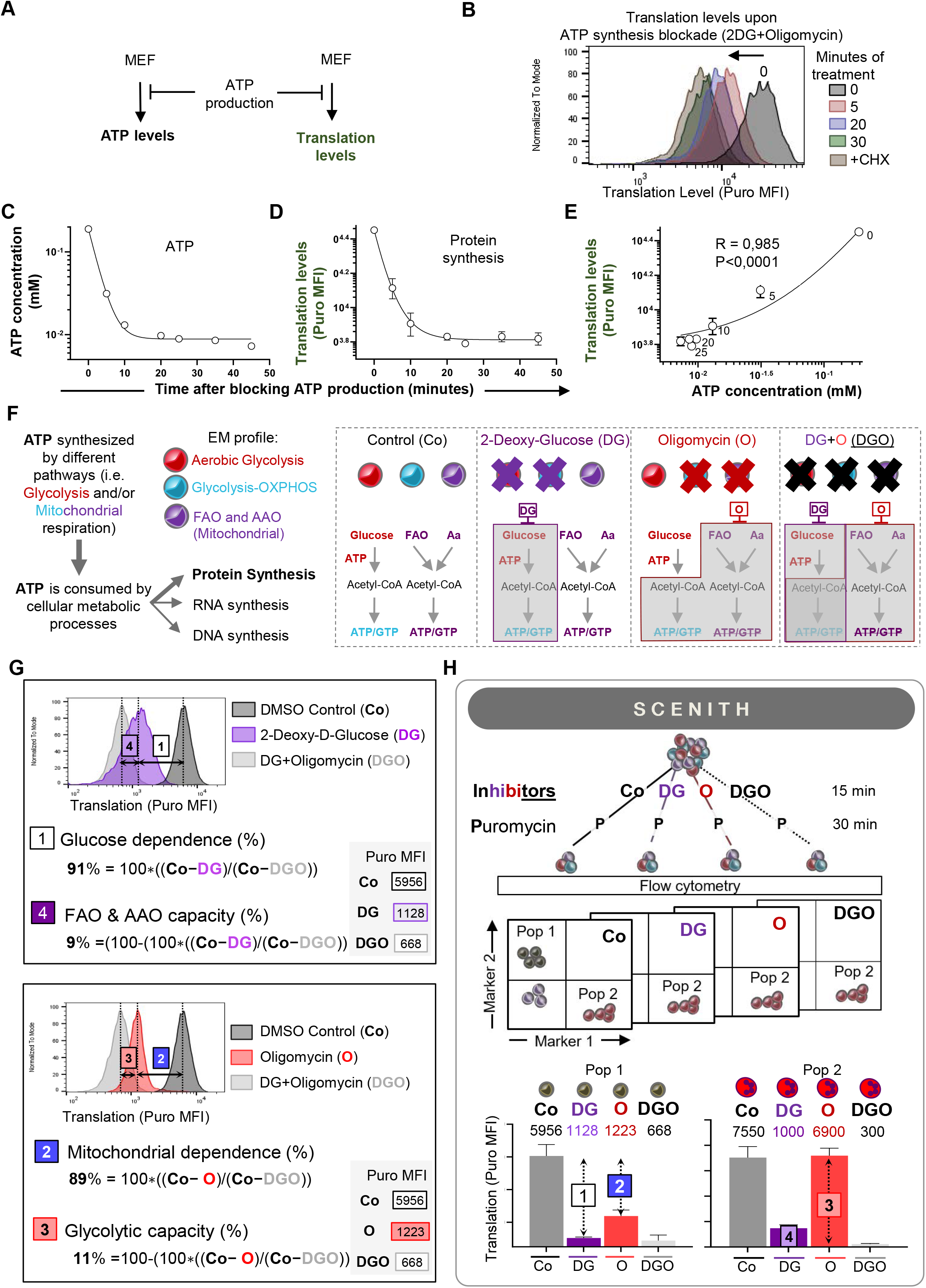
SCENITH design based on dynamic changes in protein synthesis levels upon blockade of different metabolic pathways. **(A)** Blocking ATP production and kinetics of ATP and Translation levels. **(B)** Visualization of protein synthesis after puro incorporation and staining with a new monoclonal anti-puro (clone R4743L-E8). Histogram PS level by flow cytometry in MEFs after blocking both mitochondrial respiration and glucose oxidation for different amounts of time. **(C, D and E)** Measurement in MEFs upon blocking ATP synthesis versus time of ATP levels (C), PS by flow cytometry (D) and correlation of both (E). Dot represents the means and bar the standard deviation (R=0.985, P<0.0001, N=3). **(F)** Schematic representation of a sample that contains three cell types with different metabolism profiles (Aerobic glycolysis, Glycolysis/OXPHOS, FAO and AAO/OXPHOS). Treating the mix of cells with specific drugs (DG or O) will affect each cell subset in a different way. **(G)** Examples of metabolic monitoring using SCENITH. The glucose dependence and FAO and AAO capacity; and the mitochondrial dependency and glycolytic capacity can be calculated from the MFI of puro in the different treatments following the formulas (see materials and methods). **(H)** Description of SCENITH procedure. Extract the sample, divide it and treat each with the inhibitors (e.g. DG, O, DG+O, H) and puro. After staining and flow cytometry, the profile of response of the different cells subsets is analyzed. The profile reveals the metabolic capacities and dependencies of the cells (i.e. high glucose dependence “pop 1”; and high glycolytic capacity profile “pop 2”).

As both ATP and PS levels are kinetically coupled, puromycin fluorescence (measuring PS) can act as a surrogate for energetic status in the context of flow cytometry. The principle of SCENITH is thus to incubate a given sample in parallel with specific inhibitors of known EM pathways. If a cell population is energetically dependent on a particular pathway, its ATP content will immediately drop, and so will its protein synthesis levels. The latter will be established using puromycin incorporation and its detection with fluorescent anti-puromycin antibody (Fig. 1f and Supplementary fig. 1b). SCENITH allows EM profiles to be measured in heterogenous cell populations at single cell resolution by combining cell identification and puromycilation detection by multiparametric flow cytometry (FCM) (Fig. 1g). Using PS levels, the level of glucose dependence (Gluc. Dep.) can be calculated to quantify the proportion of the PS, and therefore of ATP/GTP production, dependent on glucose oxidation (Fig. 1f, see Materials and Methods). The mitochondrial dependence (Mitoc. Dep), namely the proportion of PS dependent on oxidative phosphorylation, can be similarly established. Two additional derived parameters, “Glycolytic capacity” (Glyc. Cap.) and “Fatty acids and amino acids oxidation capacity” (FAAO Cap.) can also be calculated. Glycolytic capacity is defined as the maximum capacity to sustain PS when mitochondrial OXPHOS is inhibited (Fig. 1f, see statistic in Materials and Methods section). Conversely, FAAO Capacity is defined as the capacity to use fatty acids and aminoacids as sources for ATP production in the mitochondria, when glucose oxidation is inhibited (Glycolysis and Glucose derived Acetyl-CoA by OXPHOS) (Fig. 1f and Supplementary fig. 1b).

### SCENITH recapitulates Seahorse^®^ EM profiling of steady state and activated T cells

The metabolic switch of T cells to aerobic glycolysis upon activation was originally documented in the 1970s(Roos and Loos, 1973) and more recently confirmed using the Seahorse^®^ technology (van der Windt et al., 2012; Van Der Windt et al., 2013). To benchmark our method, we monitored the variations in EM observed in isolated bulk human blood T cells at steady state or upon activation by Seahorse^®^ in parallel to SCENITH (Fig. 2a). Upon activation, an increase in the glycolytic capacity of T cells was measured with both methods (Fig. 2b and 2c, respectively) in excellent agreement (correlation Spearman r squared 0.85, P<0.01) (Fig. 2d). We observed a significant decrease in the spare respiratory capacity in bulk T cells upon activation with Seahorse^®^ (Supplementary fig. 2a and 2b). Interestingly, an increase in oxygen consumption rate (OCR) by Seahorse^®^, was paralleled with a strong increase in the global level of PS measured by SCENITH although to a larger extent (Fig. 2e and 2f, respectively). Overall, the EM profiles of T cells upon activation obtained by Seahorse^®^ and by SCENITH were therefore very consistent. The level of translation (Fig. 2f) correlated with the global metabolic activity of the cells, and changes in the response to inhibitors confirmed the metabolic switch towards aerobic glycolysis that occurs upon T cell activation. However, SCENITH showed two main advantages over Seahorse^®^ measurements. First, the magnitude of the change in the glycolytic capacity and the standard error of the measurements with SCENITH were superior (Fig. 1b vs 1c, 1e vs 1f). Second, SCENITH analysis was performed with 10 fold less T cells (1,2.10^5^ in triplicates vs. 1,2.10^6^ cells, respectively). Moreover, SCENITH could incorporate a full spectrum of T cell markers in the analysis allowing to study in parallel the different CD3^+^ T cells subpopulations present into the bulk sample (Fig. 2), encompassing naïve, memory and effector CD4^+^ or CD8^+^ T cells subsets.

**Figure 2.**
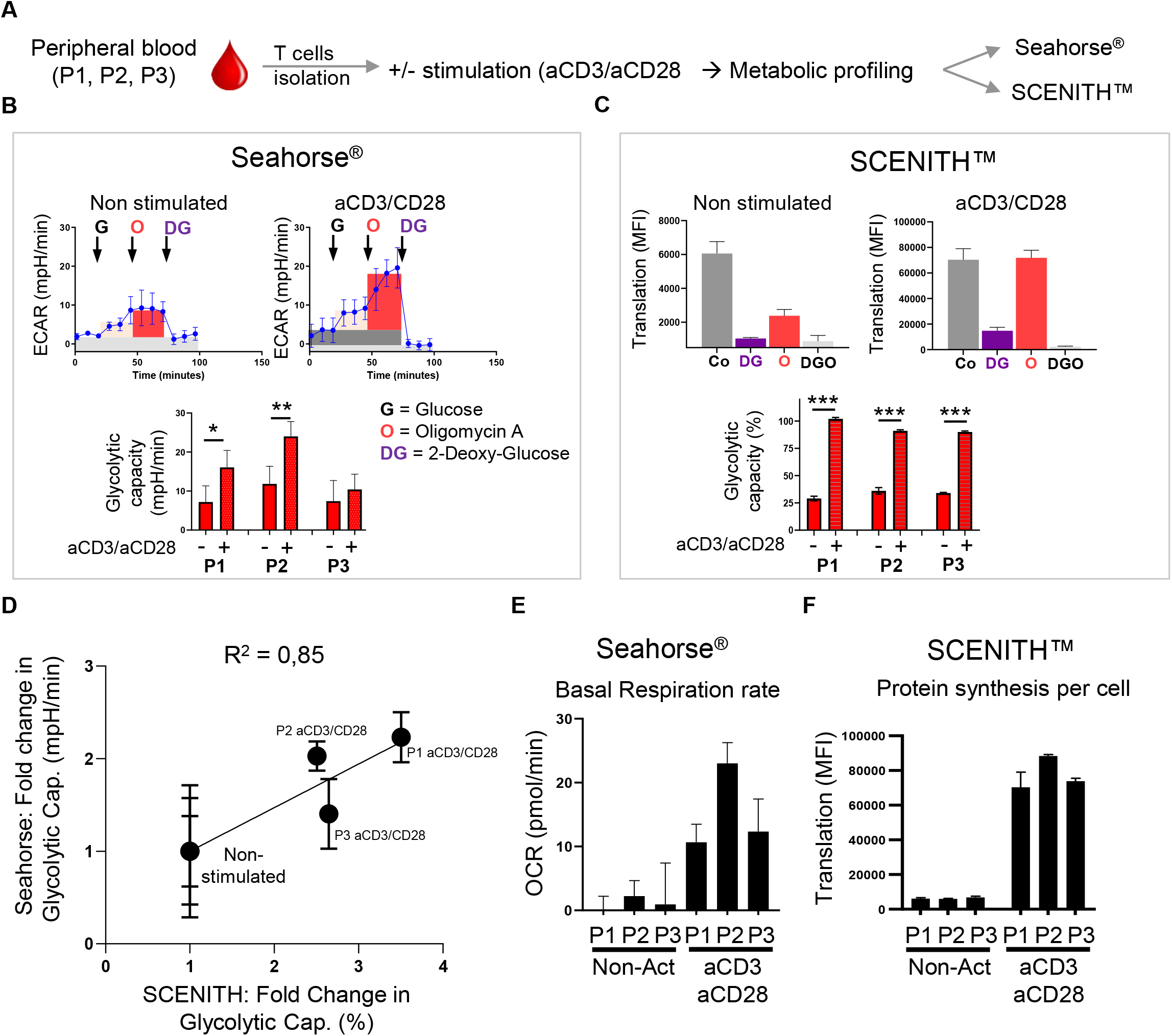
Parallel Seahorse and SCENITH metabolic analysis of resting and activated T cells. **(A)** Scheme of the experiment for analysis of resting and activated T cells. **(B and C)** Metabolic profile of T cells from three healthy donors (P1, P2, P3) analyzed with Seahorse (B) and SCENITH (C). ECAR and translation levels of non-activated and activated T cells (P1) and glycolytic capacity from both method is shown (*P<0.05; **P<0.01, ***P<0.001, N=3 each in triplicates). **(D)** Correlation between the changes in glycolytic capacity of steady state and activated T cells from three donors measured by Seahorse and SCENITH (Pearson r=0,92; R^2^=0,85; P<0.01, N=3). **(E)** Basal Oxygen Consumption Rate (OCR) in non-activated (non-Act) and activated T cells. Each bar represents the mean of P1, P2, and P3 (in triplicates). **(F)** Basal translation levels (anti-Puro gMFI) in non-activated (non-Act) and activated T cells (aCD3/CD28). Bars represent the mean of P1, P2, and P3. **(G)** SCENITH metabolic profile of whole blood directly treated with inhibitors with or without pre-incubation (1:4 V/V) in DMEM 10% FCS during 3hs. Data represents pooled whole blood from three mice (in duplicates) from three independent experiments. Two-way ANOVA, multiple comparisons.

### Metabolic deconvolution of blood T cell subsets by SCENITH identifies a memory CD8^+^ T cells subset constitutively displaying high glycolytic capacity

To expand upon the ability to deconvolve T cell subpopulations, we next applied SCENITH to mixed populations that due to heterogeneity are inaccessible to EM monitoring by Seahorse^®^. We took advantage of CD45RA, IL7RA (CD127), CCR7, CD45RO, CD57, PD1, and Perforin expression to identify and analyse the naïve and memory CD4+ and CD8+ T cell subsets present in total human blood draws. Immunodetection of these nine markers yielded six phenotypically distinct clusters/subpopulations with different abundances (Fig. 3a and 3b). The metabolic profiles of non-activated naïve T cells (CD8 or CD4), as well as memory (EM and CM) CD4 and highly differentiated CD8 (HDE) showed a medium-high degree of mitochondrial dependence (Fig. 3c), consistet with what was previously reported on their metabolic activity(Pearce et al., 2009). In contrast, the less abundant cell subsets such as CD8 early effector memory (EEM) and Natural Killer (NK) cells (that co-purfied with T cells and represented just 5% of the cells) showed higher glycolytic capacity. To determine if similar metabolic trends are observed in other species and preparations, we performed SCENITH on resting and activated mouse splenic T cells (Supplementary fig. 3a and 3b) and human blood central memory CD4 T cell subsets (Supplementary fig. 3c), demonstrating a consistent switch towards high glycolytic capacity and high glucose dependence in both mouse and human T cells upon activation (Supplementary fig. 3). During bulk analysis, naïve cells represented 42%, (the majority out of 78%) of the T cells and thus likely dominate the EM monitoring performed by Seahorse^®^ (Fig. 2e and Fig. 3c). Consequently Seahorse^®^ measurements indicate a rather low “mean” glycolytic rate/capacity and high “mean” oxygen consumtion rate (Fig. 2b) and is thus in accordance with the metabolism of naïve T cells determined by SCENITH (Fig. 3c, 3d). However, the presence of CD8^+^ EEM that display high glycolytic capacity, but represent no more that 5% of the T cells present in the sample (2000 cells) remained completely masked during Seahorse^®^ analysis.

**Figure 3.**
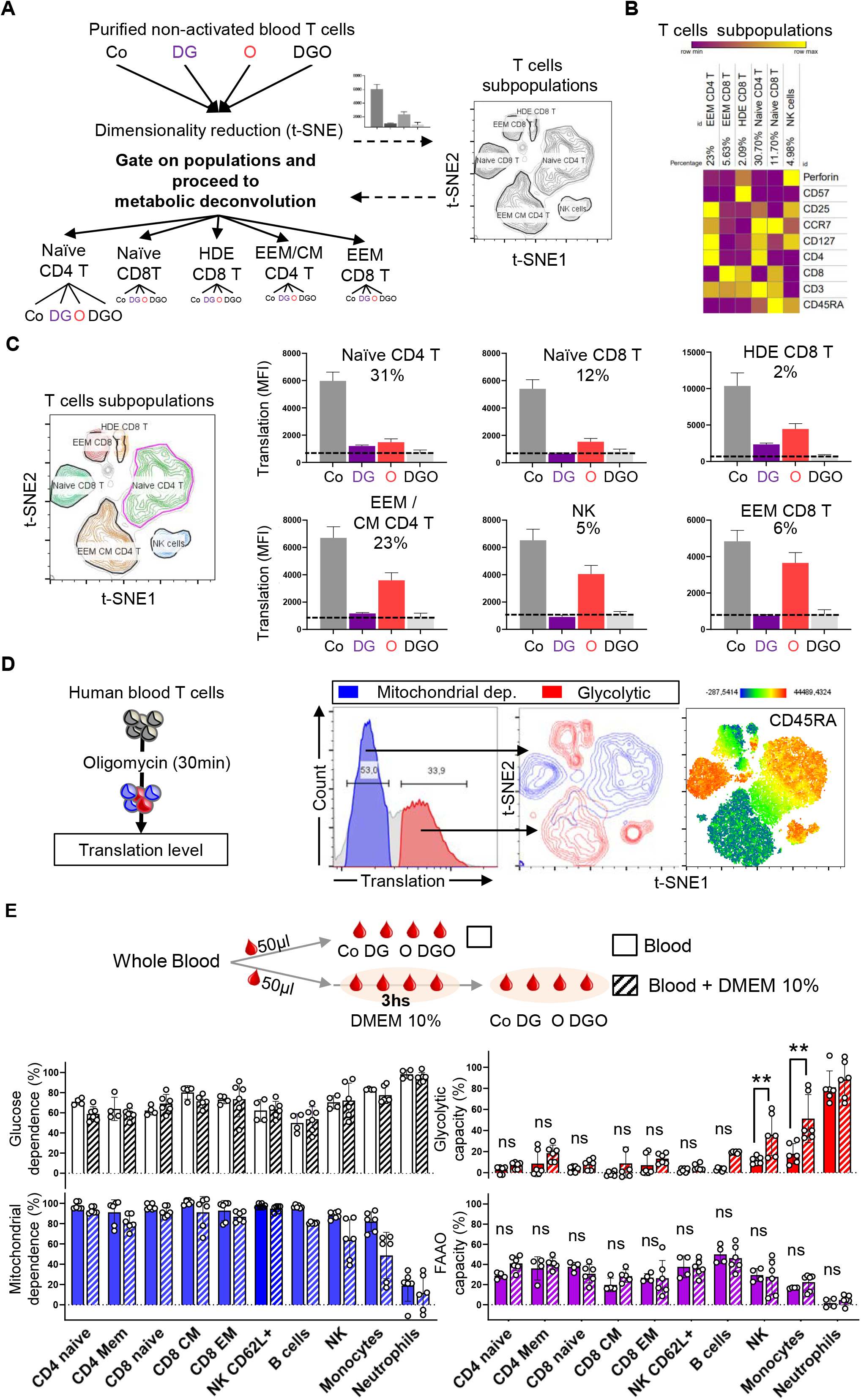
Metabolism profile of resting human blood T cells by SCENITH identifies different metabolic profile of human T cells subsets. **(A)** SCENITH analysis pipeline of T cells purified from human blood (95% pure). Dimensionality reduction (t-SNE) based on phenotypic markers is performed to the concatenated treated cells (Co, DG, O, and DGO). **(B)** Heatmap showing the level of expression of each marker (gMFI) in each cluster/subset from the t-SNE after dimensionality reduction. **(C)** Metabolic profile of the T cells subsets identified (Naïve CD4 and CD8 T cells in green, memory CD4 and HDE CD8 T cells in orange, EEM CD8 T cells in red and NK cells in blue) after SCENITH analysis. Representative translation level (anti-Puro gMFI) (P1) is shown (N=3). Black line represents background level obtained after DG+O treatment. **(D)** Two distinct metabolic profiles in human blood T cells after O treatment (left panel) revealing glycolytic and mitochondrial dependent T cells subsets. Histogram show the level of translation in all T cells (light grey line) upon mitochondrial inhibition, indicating the presence of “glycolytic” cells subsets (in red) and “mitochondrial dependent” cells (in blue). Gating them into the t-SNE plot (right panel) to identify the phenotype of “glycolytic” and “mitochondrial dependent” cells (blue). The marker of antigen experience CD45RA, lost in cells that have been previously exposed to TCR stimulations correlates with the metabolic profile. **(E)** Metabolic changes induced by short-term incubation of blood with cell culture media. Metabolic parameters of cell types when blood is pre-incubated with DMEM 10% FCS (0, or 3hs) or directly incubated with the inhibitors (i.e. Co, DG, O, DGO or Harringtonine) and puro. Data from pooled whole blood from three mice (in duplicates) from three independent experiments is shown (N=3). Statistical significance between both conditions T-test (** p<0.005).

Another important feature of the multiparametric SCENITH resolutive power is the possibility to feature single cell behaviors according to their sensitivity to metabolic inhibitors independently of their phenotype. This allows to identify functional metabolic heterogeneity first, and to determine the phenotype or sort different cells afterwards. As a proof of concept, resting purified T cells were treated with Oligomycin to inhibit mitochondrial respiration prior to translation monitoring. The histogram plotting translation levels showed two T cell subpopulations, one with high and one with low levels of translation (Fig. 3d). The population that showed high level of translation upon mitochondrial inhibition were labeled as “Glycolytic” (Fig. 3d, in red) and the cells with blocked translation as “Mitochondrial dependent” (in, blue). As shown using t-distributed stochastic neighbor embedding (t-SNE), the phenotype of Glycolytic and Respiratory T cells recapitulated our previous results (Fig. 3a-c) and showed that the expression of CD45RA, mostly present in naïve T and NK cells, correlated well with “Mitochondrial dependence” (Fig. 3d). In conclusion, we found that SCENITH allows for both the measurement of the EM profile of known non-abundant cell subsets of interest, but also the sorting and identification of “unknown” cells with specific metabolic profile present within an heterogenous sample.

### Metabolic profiling of mouse and human myeloid cell subsets

Compared to T cell subsets, the metabolism profile of myeloid cell subsets from human and mouse tissue origins has been far less studied(Saha et al., 2017). Among myeloid cell subsets, dendritic cells (i.e. DC1, DC2, DC3 and pDCs) are non-abundant professional antigen presenting cells (APCs) which serve as sentinels for the immune system. Each subset expresses a particular set of microbial pattern recognition receptors and are specialized in activation of CD8^+^ (i.e. DC1), CD4^+^ (i.e. DC2 and DC3) T cells and antiviral cytokine production (pDCs). For instance, Lipopolysaccharide (LPS) detection by TLR4 on DC2 and DC3 promotes activation by triggering a series of signaling cascades resulting in changes in gene expression, in membrane traffic and in energetic metabolism(Amiel et al., 2012; Everts et al., 2012; Krawczyk et al., 2010). Dendritic cells patrol all tissues emanating from the blood stream where they represent a very small fraction of the PBMC, making the isolation of millions of DC1, DC2 or pDC from the same donor very challenging. We therefore decided to use SCENITH to perform EM profiling of blood myeloid cell subsets from heathly donors as well as of mouse bone marrow- and spleen-derived DCs stimulated or not to generate a detailed metabolic atlas of the myeloid cell populations present in mouse or human blood^24^.

Following deconvolution, we first ranked myeloid cells by glucose dependency, finding that classical monocytes (Mono1, CD14^+^CD16^−^) were the most glucose dependent, whereas the DC precursors (DC5) were the least (Fig. 4a). These populations then lay near the extremes of mitochondrial dependence, where DC5 display the highest and Mono1 the lowest. However, we also observed examples of cell subsets (e.g. pDC and Mono2) that were highly glucose dependent as well as mitochondrial dependent and had only moderate glycolytic capacity. In contrast, some (DC1 and DC2) displayed relatively high glycolytic capacity and moderate glucose dependency, suggesting a certain degree of metabolic plasticity. Cells ranked in opposite order for FAAO capacity compared to glucose dependency, consistent with the idea that cells with low glucose dependence can sustain translation and energy production by free-fatty and amino acids oxidation.

**Figure 4.**
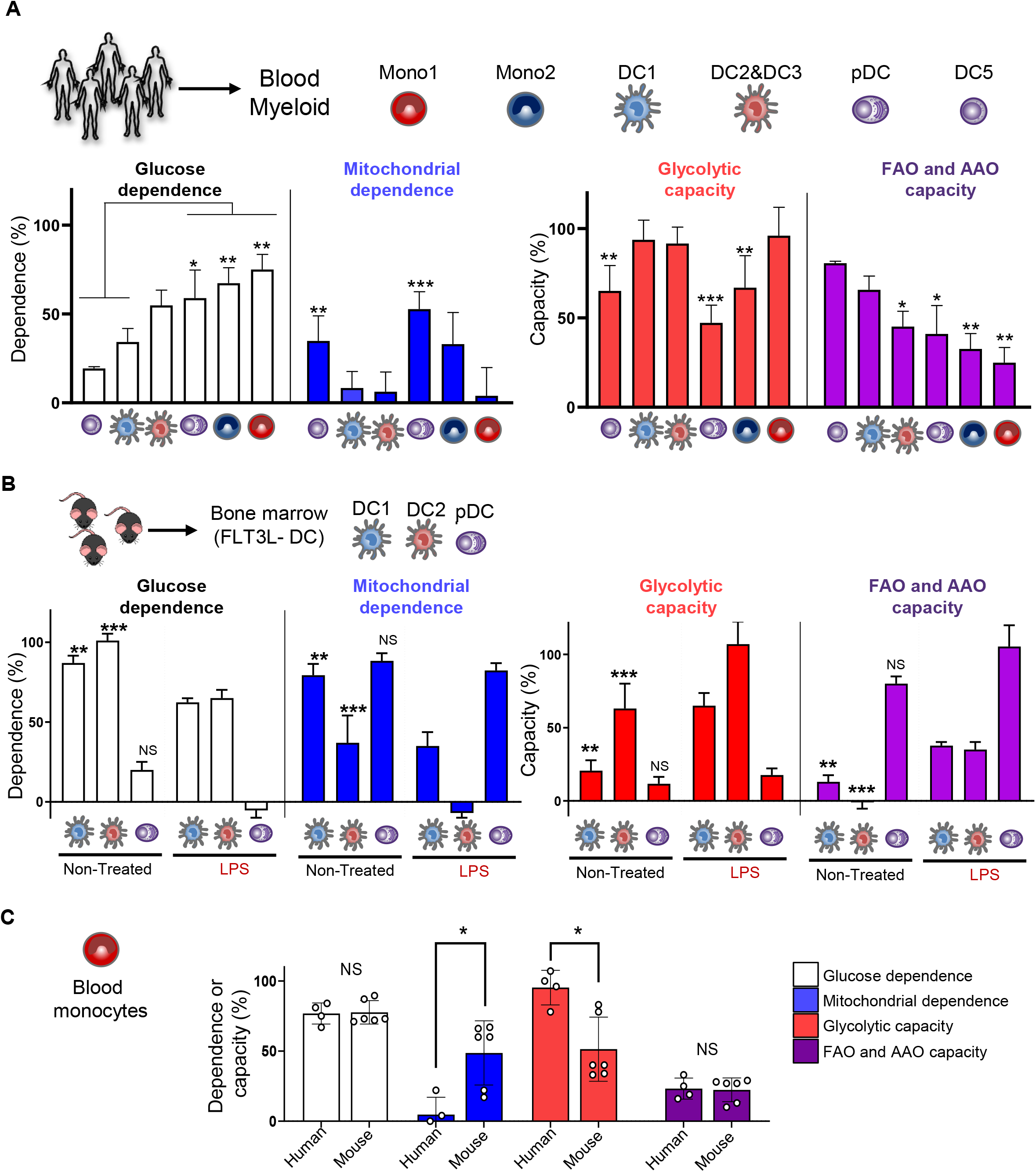
Metabolic profile of human blood DCs and monocytes, and mouse bone marrow derived DCs using SCENITH. **(A)** Metabolic profile of human blood monocytes and DC subsets obtained by SCENITH. (N=5 independent healthy donors). Statistical significance two-way ANOVA comparing all columns was performed (* p<0.05; ** p<0.005; ****p<0.0001). The pDC, the Mono2 or Mono1, showed statistically significant differences against DC1 or DC5. **(B)** Metabolic profile of mouse bone marrow derived DCs (FLT3L-DC) obtained by SCENITH (N=3) in non-treated vs LPS treated cells two-way ANOVA * p<0.05; ** p<0.005; ****p<0.0001. **(C)** Metabolic profile of blood monocytes from human (N=4) or mouse (N=9, 3 mice pooled, in duplicates, three independent experiments). Statistical significance between human and mouse monocytes by T-test (* p<0.05).

To test whether a rapid metabolic switch occurs upon TLR4 activation in human myeloid cells, we used an antibody panel for analyzing Mono1, Mono2, DC1, DC2 and pDCs in PBMCs treated with LPS. Most subsets did change their metabolic profile, while Mono1 maintained their glycolytic profile but increased their global level of translation (Supplementary fig. 4a). The only subset that changed its metabolic profile were the DC1 that showed a mild increase in glucose dependence and a moderate decrease in mitochondrial dependence (Supplementary fig. 4a).

To gain insight into the metabolic profile of mouse DC populations, we analyzed the metabolism profile of mouse bone marrow-derived DC1, DC2 and pDC (FLT3L-BMDC) at steady state and upon activation with LPS (Fig. 4b). In accordance to what was observed in steady state human blood DCs, mouse DC2 subset showed the highest glycolytic capacity, followed by DC1 and pDC. While Glucose dependence was relatively high in DC1 and DC2 from human and mouse (40-55% vs. 80-100%, respectively), an important difference was observed in the glucose dependency of mouse versus human pDCs (15% vs 60%). LPS treatment shifted EM profile towards lower mitochondrial dependence of DC1 and DC2. However only DC2, showed an increased global level of protein synthesis, probably reflecting the abundant TLR4 expression in this DC subset‥ Considering that we analyzed DC isolated from human blood and derived in-vitro from mouse bone marrow, SCENITH EM profiles are surprisingly consistant. These results confirm that SCENITH allows to identify different cell populations sharing similar EM profile and that EM in DCs varies according to their state of activation.

### Profiling the metabolic state of human tumor-associated myeloid cells

Immunotherapies are a game changer in oncology yet only a fraction of patients shows complete immune-mediated rejection of the tumor. The variations observed in patients responses to treatment have created a strong need for understanding the functional state of tumor-associated immune cells (immunoprofiling)(Galon et al., 2006). We thus used SCENITH to perform paralleled phenotypic and metabolic profiling of human tumor samples, and investigate the heterogeity of immune cell subsets by comparing tumors of diverse origins with tumor-free adjacent tissue. We analyzed PBMCs from healthy donors, explanted meningioma, brain metastasis (originated from a breast cancer), as well as renal carcinoma tumors and renal juxtatumoral tissue. In the case of renal cancer, both SCENITH and single cell RNA seq analysis were performed in parallel on the same sample.

We observed 8 different myeloid populations in meningioma and 6 different subset in renal carcinoma (Fig. 5a and 5b, respectively; Supplementary fig. 5a and 5b)), that were all profiled by SCENITH (Fig. 5c). Upon clustering of the different cell subsets based on EM profiling, two groups emerged, a “Glycolytic cluster” and a “Respiratory cluster” (Fig. 5c and Supplementary fig. 5c). Mono1 and Neutrophils displayed glycolytic metabolism profiles in all blood samples and tumors tested (Fig. 5d). In contrast, Mono2, DC1 and DC2 showed relatively high glycolytic capacity when isolated from kidney tumor and juxtatumoral tissues, while these subsets showed high respiratory metabolism profile in the two brain tumors. Conversely, tumor-associated macrophages (TAM), showed high mitochondrial dependence, while juxta-tumoral macrophages displayed high glycolytic capacity (Fig. 5d), suggesting that tumor microenvironment modifies TAM EM. The decrease of glycolytic capacity in TAM as compared to juxta-tumoral macrophages was previously associated with increased immunosuppression in the tumor environment, tumor progression via both nutritional and immunological cues, and poor patient survival(Vitale et al., 2019). These results demonstrate again the analytical capacity and descriminative power of SCENITH, that reveals that in addition to the cancer type, the tumor anatomical origin could influence the metabolism of immune subsets, introducing an additional layer of heterogeneity in the tumor environment.

**Figure 5.**
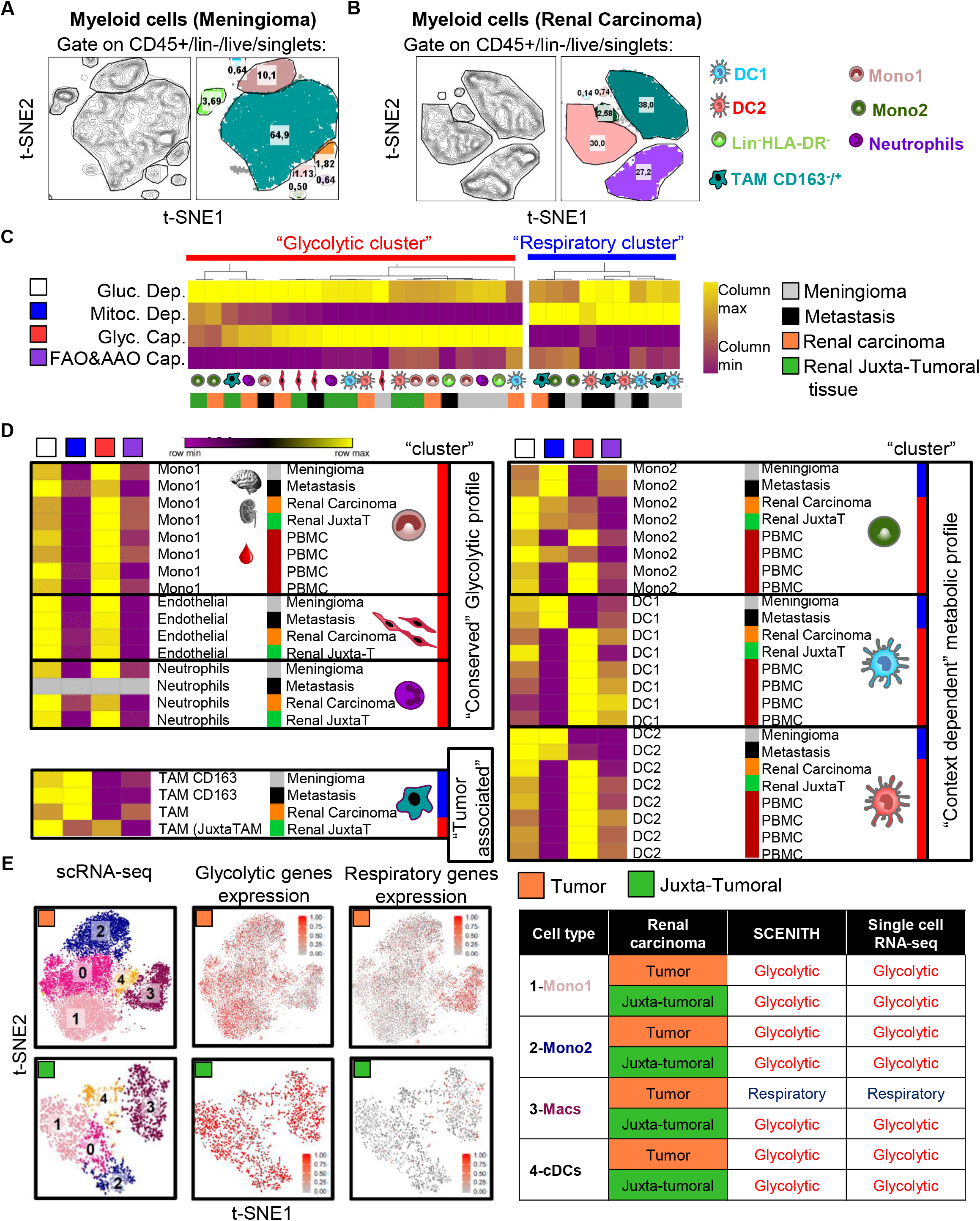
Paralleled SCENITH and scRNAseq in human tumor and juxta-tumoral samples identifies conserved metabolic profiles. **(A and B)** Myeloid subsets observed in the human meningioma tumor sample (A) and Renal carcinoma (B). Myeloid cells gated on CD45^+^/CD3^−^CD20^−^CD19^−^CD56^−^/Live-dead^−^/singlets. Number in the t-SNE represents the percentage of the population. **(C)** Heatmap of the metabolic profile (columns) of each myeloid cell subset from each type of tissue (rows). Unsupervised hierarchal clustering of subsets by metabolic profile identifies respiratory (blue bar) and glycolytic (red bar) clusters. **(D)** Reordering of the rows by cell type based on (B) to identify changes in metabolism profile in the same cell subset in the blood, the tumors and juxta-tumoral tissue. **(E)** Clusters of myeloid cells identified in the renal carcinoma and juxta-tumoral tissue by scRNAseq (left panel). Expression of glycolytic and respiratory gene signatures in all cells extracted from the tumor. Summary of the results obtained by SCENITH and scRNAseq in tumor and juxta-tumoral myeloid cells. Populations named with numbers in the t-SNE and described in table.

### Linking scRNA-seq and energetic metabolism profile in tumor-associated myeloid cells

We decided to link the descriminative power of SCENITH with single cell RNA-seq (scRNAseq) to potential clinical interest. The activation of human and mouse myeloid cells with LPS, showed that only cells expressing the LPS receptor/co-receptor (TLR4, CD14) show an increase in the global level of protein synthesis (human Mono1 and mouse DC2). In contrast, DC1 that do not respond to LPS, also showed a change in their metabolic profile, suggesting that bystander activation of DCs by Mono1 or DC2 derived cytokines can induce a change in EM profile, independently of TLR triggering.

Importantly, SCENITH can establish the metabolic profile of very scarce cells, as exemplified by the analysis of early effector T cells that represent around 5% of the total T cells isolated from blood (500 cells), thus representing a gain of sensitivity of aproximatively 800 fold compared to Seahorse^®^ measurements. This gain was even more dramatic, when the metabolism profile of DC cell subsets was established from human tumor biopsies, in which they constituted aproximately 0,5% of the myeloid cell population. These results demonstrate the analytical capacity and discriminative power of SCENITH and its potential in analysing how the anatomical and tissue context could influence energetic metabolism of immune cell subsets, thus introducing an additional layer of heterogeneity in the healthy tissues and tumor environment. Thus, SCENITH is a powerful technique that: (1) allows to rapidly establish metabolic profile of several populations of cells in an heteregenous sample by flow cytometry, (2) does not require a large amount of cells and thus avoids their expansion in culture and (3) can be combined with scRNA-seq to establish a full map of EM profile of cells. Given the direct relationship between EM and the functionality of lymphoid effector cells and myeloid cells, SCENITH analysis could be used to define the ‘immune EM contexture’ and complement the establishment of an immunoscore that’s defines immune fitness of tumours and predicts and stratifies patients for tailored therapies, aiming at manipulating metabolic pathways to improve anti-tumoral immune effector functions.

## Materials and Methods

### Cells and Cell Culture

Mouse splenocytes from WT C57BL/6J (Jackson) or PERK KO C57BL/6J background mice (Zhang et al., 2002) were cultured in DMEM containing 5% of Fetal Calf Serum (FCS) and 50 μM of 2-Mercaptoethanol (Mouse cells culture media, MCCM) at 37 °C 5% of CO2. GM-CSF BM-derived dendritic cells (GM-bmDCs) were differentiated in vitro from the bone marrow of 6–8-week-old male mice, using GM-CSF, produced by J558L cells. Bone marrow progenitors were plated at 0.8.10^6^ cells/ml, 5 ml/well in 6-well plates, and cultivated with RPMI (GIBCO), 5% FCS (Sigma-Aldrich), 20 μg/ml gentamycin (Sigma-Aldrich), 50 μM β-mercaptoethanol (VWR), and GM-CSF. The medium was replaced every 2 days; BM-derived DCs were used for experiments at day 6. Similarly, FLT3L BM-derived dendritic cells (FLT3L-bmDCs) were differentiated by adding FLT3L to RPMI, 10% of Fetal Calf Serum (FCS) and 50 μM of 2-Mercaptoethanol (Mouse cells culture media, MCCM) during 6 days at 37 °C 5% of CO2. To obtain splenocytes, eight weeks old wild type C57BL/6J mice were sacrificed by cervical dislocation and splenectomized. Single cells suspentions from the spleens were generated and cultured in MCCM as previously described. Mononuclear cell enriched from blood of healthy donors was submitted to Ficoll-paque plus (PBL Biomedical Laboratories). PBMCs and Whole blood were cultured in the absense (non stimulated) or in the presence of LPS for 4hs. Immune cell stimulations were performed in the absence (Control) or presence of 0,1 μg/ml of extrapure Lipopolysacharide (Invivogen LPS, Cat. tlrl-3pelps), 10 μg/ml Poly I:C (Invivogen, Cat. No. tlrl-pic), CpG-A ODN 2216 (Invivogen, Cat. No. tlrl-2216) or PMA (5 ng/ml; Sigma, Cat. no. P-8139) and ionomycin (500 ng/ml; Sigma, cat. no. I-0634) over night for T cell stimulations and 4 hours for Dendritic cells. T cells from different human donors (P1, P2, P3) were isolated using the RosetteSep™ negative isolation method and activated (using BD Human T cell activator beads coated with anti-CD3 and anti-CD28) or not.

### ATP measurement

2.10^4^ MEFs were seeded in 100ul of 5% FCS DMEM culture media ON in opaque 96 well plates. Cells were incubated with the inhibitors for the times indicated in the Figure. After, 100ul of Cell titer-Glo luminiscence ATP reconstituted buffer and substrate (Promega, Cat. No. G7570) was added to each well and Luminiscence was measured after 10 minutes following manufacturer instructions. A standard curve with ATP was performed using the same kit and following manufacturer instructions.

### Metabolic flux analysis (Seahorse^®^)

support our results regarding the cellular complexity of the tumor environment and correlate EM profiling with gene expression. We compared in each individual population, the functional EM profile obtained by SCENITH with a metabolic gene expression pattern obtained by scRNAseq. We first identified expression signatures of specific glycolytic and respiratory genes that correlated with the functional metabolism profiles of different blood myeloid cells (Supplementary Fig. 5d). We then tested the expression of these glycolytic and respiratory metabolic genes (mRNA levels) in the different tumor-associated myeloid populations (CD45^+^Lin^−^HLA-DR^+^). Sorted cells from the renal carcinoma and juxtatumoral tissue were subjected to scRNAseq using 10X Genomics Chromium paired with deep sequencing (Fig. 5e and Supplementary Fig. 5e). Analysis of 12,801 cells for the tumor and 2,080 for the juxta tumoral tissue yielded 6 and 5 high quality population clusters respectively. To rigourously identify the myeloid populations, we verified the expression of characteristic markers of these populations(Villani et al., 2017) and assigned cellular identities in the t-SNE representations (Supplementary Fig. 5e). We focused on 5 monocytes and macrophages clusters expressing MAFB and/or CSF1R, that were present both in tumor and juxta-tumoral tissue (Supplementary Fig. 5f). Expression of classical surface markers like FCGR3A/CD16 and CD14 (Supplementary Fig. 5g), confirmed that clusters 0 and 1 represent CD14^+^CD16^−^ classical monocytes. Cluster 2 represents CD14^−^CD16^+^ non classical monocytes (Mono2), while co-expression of CD14 and CD16 for the clusters 3 and 4 corresponds to macrophage-like populations. We performed differential expression (DE) unsupervised analysis for each of the myeloid cluster versus all the other clusters identified in the tumor bed and generated heatmaps for the top 5 most differentially expressed genes in the tumor (Supplementary Fig. 5h) and the juxta tumoral tissue (Supplementary Fig. 5h, right). In addition to highlighting key genes that contributed to the unbiased segregation of these populations in both tissue, we confirmed the high expression of macrophage specific genes by cluster 3 and 4, such as APOE, C1QC and RGS1. We next overlayed on the t-SNE plots our two EM gene signatures. Strongly correlating with SCENITH profiles, monocytes clusters (0,1,2) presented an enrichement in glycolytic genes signature both in tumor and juxta tumoral tissue. Conversely macrophages (cluster 3) showed high expression of the respiratory signature in the tumor, while, as predicted by SCENITH, this was not detectable in juxta-tumoral tissue (Fig. 5e). Monocytes-derived dendritic cells differently from macrophages, presented an enrichment in glycolytic genes signature both in tumor and juxta tumoral tissue (Supplementary Fig. 5i).

Altogether, these results indicate that tumor micro-environment specifically modifies functional EM of Macrophages by durably affecting metabolic gene expression. By correlating the results of scRNAseq analysis and SCENITH profiling on different blood myeloid cell subsets (Fig. 5f), we identify a functional gene signature (Supplementary Fig. 5d) that can be used to metabolically profile a variety of cell types and tissues, using gene expression.

## Discussion

SCENITH is a flow cytometry-based method to establish simultaneously the phenotypic and energetic metabolism profiles of multiple cell types in parallel. The method is rapid, highly sensitive and consistent with other established techniques, including Seahorse^®^. As the treatments with different inhibitory drugs are performed in parallel, SCENITH can be used to monitor cellular responses to a variety of metabolites, inhibitory compounds and their combinations. Given that flow cytometers are available in most research institutes and hospitals, SCENITH represents an accessible method to perform functional EM profiling. Compared to other methods (see Table 1), its sensitivity, accessibility, single cell resolution, stability of the readout, short experiment time, compatibility with fixation and sorting, makes of SCENITH, an unrivalled aproach for studying energy metabolism in tissues and complex populations.

The resolutive power of SCENITH highlighted important variations amongst subsets of the same cell population. During our analysis of human blood myeloid subsets, we monitored myeloid subpopulations that are known to be precursors of DC1, pDC and DC3. The precursor of DC1 and pDC, called DC5, showed an EM with intermediate characteristics between DC1 and pDC, that are both metabolically polarized. Mono1 are precursors of DC3 and even if the metabolic profile is similar, Mono1 EM is more glycolytic. Our results suggest that changes in metabolism is embedded in the myeloid differentiation program and indicate that modulating metabolism in dendritic cells precursors might influence the generation of different dendritic cells subsets with OCR and ECAR were measured with the XF24 Extracellular Flux Analyzer (Seahorse Bioscience). 4.10^5^ cells with αCD3/αCD28 beads or not, were placed in triplicates in XF medium (nonbuffered Dulbecco’s modified Eagle’s medium containing 2.5 mM glucose, 2 mM L-glutamine, and 1 mM sodium pyruvate) and monitored 25 min under basal conditions and in response to 10mM Glucose, 1 μM oligomycin, 100 mM 2-Deoxy-Glucose. Glycolytic capacity was measured by the difference between ECAR level after add oligomycin and before add glucose. OCR, ECAR and SRC parameters was analyzed and extract from Agilent Seahorse Wave Desktop software. Glycolytic capacity was obtained by the difference between ECAR level after add Oligomycin and before add Glucose.

### SCENITH

Cells were plated at 1.10^6^ cells/ml, 0,5 ml/well in 48-well plates. Experimental duplicates/triplicates were performed in all conditions. After differentiation, activation or harvesting of human of cells, wells were treated during 30 minutes with Control, 2-Deoxy-D-Glucose (DG, final concentration 100mM; Sigma-Aldrich Cat. No. D6134), Oligomycin (Oligo, final concentration 1μM; Sigma-Aldrich Cat. No.75351), or a combination of the drugs at the final concentrations before mentioned. As negative control, the translation initiation inhibitor Harringtonine was added 15 minutes before addition of Puromycin (Harringtonine, 2 μg/ml; Abcam, cat. ab141941). Puromycin (Puro, final concentration 10 μg/ml; Sigma-Aldrich, Cat. No. P7255) is added during the last 15 minutes of the metabolic inhibitors treatment. After puro treatment, cells were washed in cold PBS and stained with a combination of Fc receptors blockade and fluorescent cell viability marker, then primary conjugated antibodies against different surface markers (see FACS material and methods) during 30 minutes at 4°C in PBS 1X 5% FCS, 2mM EDTA (FACS wash buffer). After washing with FACS wash buffer, cells were fixed and permeabilized using BD Cytofix/Cytoperm™ (Catalog No. 554714) following manufacturer instructions. Intracellular staining of Puro using our fluorescently labeled anti-Puro monoclonal antibody was performed by incubating cells during 1 hour (1:800, Clone R4743L-E8, in house produced and conjugated with Alexa Fluor 647, to obtain 10 times better signal to noise ratio than commercially available monoclonal antibodies) at 4°C diluted in Permwash. For SCENITH troubleshooting see Table 2.

### Patients and samples

The renal carcinoma patient enrolled in this study provided written and informed consent to tissue collection under a University of California, San Francisco (UCSF) institutional review board (IRB)-approved protocol (UCSF Committee on Human Research (CHR) no. 13-12246). The meningioma and brain meastasis patients enroled in this study provided written and informed consent in accordance with institutional, national guidelines and the Declaration of Helsinki. This protocol was approved by institutional review board (AP-HM CRB-TBM tumor bank: authorization number AC-2018-31053, B-0033-00097).

### Processing of and mouse solid tumors SCENITH

0,2-0,4 grams of solid tumor tissue was partially dissociated using chirurgical scissors or tissue chopper (McIlwain Tissue Chopper^®^ Standard plate) to generate “tumor explant suspention”. Tissue explants suspention, containing tissue cubes of approximately 400μm of cross section, were put in suspention in complete RPMI media and incubated directly with control or metabolic inhibitors, and with Puromycin following the SCENITH protocol. Next, tumor explants were dissociated using Tissue Liberase and DNAseI with the help of a Gentle Macs (Miltenyi) following manufacturers instructions. Cell suspentions were washed, counted and 2-5.10^6^ total cells were seed in triplicates before proceeding with lived dead and FC block staining. Next, cells were stained for surface makers, fixed and permeabilized (ThermoFisher, FOXP3 fixation kit) and stained for nuclear and cytoplasmic markers as mentioned above.

### Human single cell RNA-sequencing

Live CD3-CD19/20-CD56-cells were sorted from renal carcinoma tumor and juxta tumoral tissue using a BD FACSAria Fusion. After sorting, cells were pelleted and resuspended at 1.10^3^ cells/μl in 0.04%BSA/PBA and loaded onto the Chromium Controller (10X Genomics). Samples were processed for single-cell encapsulation and cDNA library generation using the Chromium Single Cell 3’ v2 Reagent Kits (10X Genomics). The library was subsequently sequenced on an Illumina HiSeq 4000 (Illumina).

### Single cell data processing

Sequencing data was processed using 10X Genomics Cell Ranger V1.2 pipeline. The Cell Ranger subroutine mkfastq converted raw, Illumina bcl files to fastqs which were then passed to Cell Ranger’s count, which aligned all reads using the aligner STAR (Dobin et al., 2013)ref against GRCh38 genomes for human cells. After filtering reads with redundant unique molecular identifiers (UMI), count generated a final gene-cellular barcode matrix. Both mkfastq and count were run with default parameters.

### Cellular Identification and Clustering

For each sample, the gene - barcode matrix was passed to the R (v. 3.6.0) software package Seurat(Satija et al., 2015) (http://satijalab.org/seurat) (v3.1.1) for all downstream analyses. We then filtered on cells that expressed a minimum of 200 genes and required that all genes be expressed in at least 3 cells. We also removed cells that contained > 5% reads associated with cell cycle genes(Kowalczyk et al., 2015; Macosko et al., 2015). Count data was then log2 transformed and scaled using each cell’s proportion of cell cycle genes as a nuisance factor (implemented in Seurat’s ScaleData function) to correct for any remaining cell cycle effect in downstream clustering and differential expression analyses. For each sample, principal component (PC) analysis was performed on a set of highly variable genes defined by Seurat’s FindVariableGenes function. Genes associated with the resulting PCs (chosen by visual inspection of scree plots) were then used for graph-based cluster identification and subsequent dimensionality reduction using t-distributed stochastic neighbor embedding (tSNE). Cluster-based marker identification and differential expression were performed using Seurat’s FindAllMarkers for all between-cluster comparisons.

### Flow cytometry

Flow cytometry was conducted using BD Symphony and BD LSR Fortessa X-20 machine (BD Biosciences™) and data were analyzed with FlowJo (Tree Star™) or FLOWR-software (Guillaume VOISSINE, https://github.com/VoisinneG/flowR). The antibodies used to stain mouse splenocytes were anti-Puro-Clone R4743L-E8, rat IgG2A in house produced and (conjugated with Alexa Fluor 647 or Alexa-Fluor 488), anti-Ki67 PE-eFluor-610 (eBioscience™, Catalog No. 61-5698-82) CD4-APC-eF780 (eBioscience™, Catalog No. 47-0042-82), CD8-APC (eBioscience™, Catalog No. 17-0081-83), CD80-PercPCy5.5 (Biolegend™, Catalog No. 104722), anti-B220-BV421 (Biolegend™, Catalog No. 103251), anti-MHC-II-AF700 (eBioscience™, Catalog No. 56-5321-82), LIVE/DEAD™ Fixable Aqua Dead Cell Stain (Invitrogen™, Catalog No. L34957). The following anti-Human antigens antibodies were used for staining whole blood and PBMCs upon SCENITH protocol application. Alexa Fluor-488 Mouse Anti-Human Axl (Clone 108724, R&D Biosystems, Cat. No. FAB154G), BUV395 Mouse Anti-Human CD11c (Clone B-ly6, BD Bioscience, Cat. No. 563787), BUV737 Mouse Anti-Human CD86 (Clone FUN-1, BD Bioscience, Cat. No. 564428), BV510 Mouse Anti-Human CD19 (Clone HIB19, BD Bioscience, Cat. No. 740164), BV510 Mouse Anti-Human CD3 (Clone HIT3a, BD Bioscience, Cat. No. 564713), BV510 Mouse Anti-Human CD56 (Clone B159, BD Bioscience, Cat. No. 740171), BV605 Anti-Human HLA-DR (Clone L243, BioLegend, Cat. No. BLE307640), BV650 Mouse Anti-Human CD16 (Clone 3G8, BD Bioscience, Cat. No. 563692), BV711 Mouse Anti-Human CD14 (Clone M5E2, BD bioscience, Cat. No. 740773), BV785 Mouse Anti-Human CD45RA (Clone HI100, BioLegend, Cat. No. BLE304140), Live Dead Fixable Aqua Dead Cell Stain Kit (Life Technologies, Cat. No. L34957), PE Rat Anti-Human Clec9A/CD370 (Clone 3A4, BD Bioscience, Cat. No. 563488), PE-Cy7 Mouse Anti-Human CD22 (Clone HIB22, BD Bioscience, Cat. No. 563941), AF488 Mouse Anti-Human CD38 (Clone HIT-2, BioLegend, Cat. No. BLE303512).

### Animal studies

Wild type C57BL/6 mice were purchased from Charles River and maintained in the animal facility of CIML under specific pathogen-free conditions. This study was carried out in strict accordance with the recommendations in the Guide for the Care and Use of Laboratory Animals the French Ministry of Agriculture and of the European Union. Animals were housed in the CIML animal facilities accredited by the French Ministry of Agriculture to perform experiments on alive mice. All animal experiments were approved by Direction Départementale des Services Vétérinaires des Bouches du Rhône (Approval number A13-543). All efforts were made to minimize animal suffering.

### Statistical analysis

Statistical analysis was performed with GraphPad Prism software. When several conditions were to compare, we performed a one-way ANOVA, followed by Tukey range test to assess the significance among pairs of conditions. When only two conditions were to test, we performed Student’s t-test or Welch t-test, according the validity of homoscedasticity hypothesis (* P<0.05, ** P<0.01, *** P<0.005).

### Calculation and meaning of SCENITH derived parameters

To quantify the different energetic metabolism parameters that constitute the metabolic profile of a cell, such as pathways dependency, we used simple algorithms that quantifiy the relative impact of inhibiting a given pathway compared to a complete inhibition of ATP synthesis (Fig. 1f). While SCENITH allow the use of any combination of metabolic or signalling inhibitors, herein we focused on inhibitors of glycolysis and of mitochondrial respiration to derive metabolic parameters. The percentual of glucose dependence (Gluc. Dep.) quantifies how much the translation levels are dependent on glucose oxidation. Gluc. Dep. is calculated as the difference between PS levels in 2-Deoxy-D-Glucose (DG) treated cells compared to control (Co), divided by the difference in PS upon complete inhibition of ATP synthesis (DG, FCCP and Oligomycin A, combined; treatment Z) compared to control cells (Fig. 1f). In a similar fashion, percentual mitochondrial dependence (Mitoc. Dep) quantifies how much translation is dependent on oxydative phosphorylation. Mitoc. Dep. is defined as the difference in PS levels in Oligomycin A (“O”, mitochondrial inhibitor) treated cells compared to control relative to the decreased in PS levels upon full inhibition of ATP synthesis inhibition (treatment Z) also compared to control cells (Fig. 1f). Two additional derived parameters, “Glycolytic capacity” (Glyc. Cap.) and “Fatty acids and amino acids oxidation capacity” (FAAO Cap.) were also calculated. Glycolytic capacity is defined as the maximum capacity to sustain protein synthesis levels when mitochondrial OXPHOS is inhibited (Fig. 1f, see statistic in Materials and Methods section). Converserly, FAAO Capacity is defined as the capacity to use fatty acids and aminoacids as sources for ATP production in the mitochondria when glucose oxidation is inhibited (Glycolysis and Glucose derived Acetyl-CoA by OXPHOS) (Fig. 1f and Supplementary fig. 1b). While the total level of translation correlates with the global metabolic activity of the cells, the dependency parameters underline essential cellular pathways that cannot be compensated, while “capacity”; as the inverse of dependency, shows the maximun compensatory capacity of a subpopulation of cells to exploit alternative pathway/s when a particular one is inhibited (Fig. 1f and Supplemenary Fig. 1c).

For standard deviation calculation of SCENITH, we followed the propagation of error that is required when the means of means are used into a formula:

For error calculation:

*Co= GeoMFI of anti-Puromycin-Fluorochrome upon Control treatment*
*DG= GeoMFI of anti-Puromycin-Fluorochrome upon 2-Deoxy-D-Glucose treatment*
*O= GeoMFI of anti-Puromycin-Fluorochrome upon Oligomycin A treatment*
*Z= GeoMFI of anti-Puromycin-Fluorochrome upon DG+O treatment*

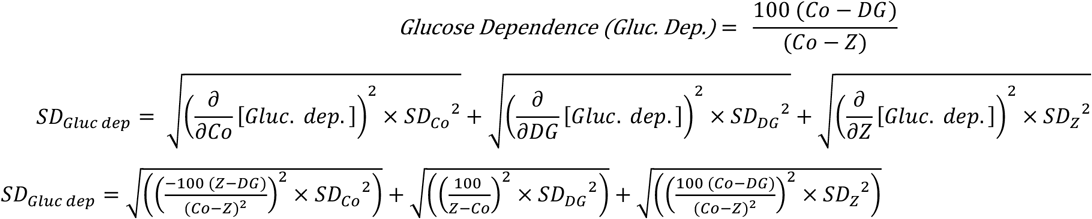

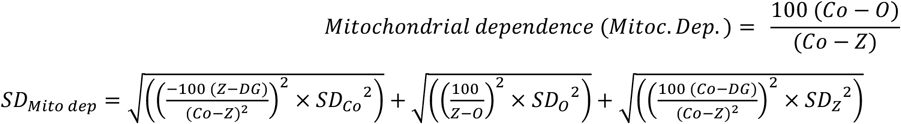

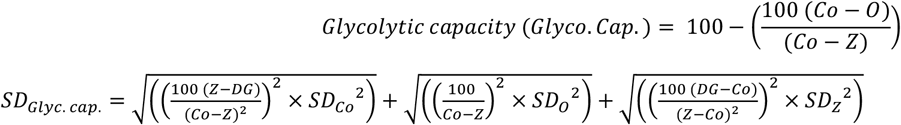

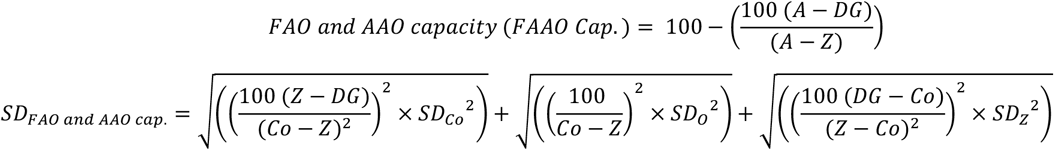

## Acknowledgments

The PP laboratory is “Equipe de la Fondation de la Recherche Médicale” (FRM) sponsored by the grant DEQ20140329536. The project was supported by grants from l’Agence Nationale de la Recherche (ANR), « ANR-FCT 12-ISV3-0002-01», A*MIDEX project “CSI” (ANR-11-IDEX-0001-02), «DCBIOL Labex ANR-11-LABEX-0043 », « INFORM Labex ANR-11-LABEX-0054 » funded by the “Investissements d'Avenir” French government program and initiated through Contrats Proof of concept CoPoc 2019 INSERM “ZIPT R19011AS” to RJA, Bourse Mobilité Cancéropôle (to UCSF) to RJA and Cancéropôle PACA "Emergence" grant “HiDi-Glio” to RJA and Projet Fondation ARC 2016 to PP. The research was also supported by the Ilídio Pinho foundation and FCT - Fundação para a Ciência e a Tecnologia - and Programa Operacional Competitividade e Internacionalização - Compete2020 (FEDER) – references POCI-01-0145-FEDER-016768 and POCI-01-0145-FEDER-030882. We acknowledge financial support from n° ANR-10-INBS-04-01 France Bio Imaging and the ImagImm CIML imaging. We thank Noella LOPES-PAPPALARDO and Alexandre BOISSONNAS for reading the manuscript and suggestions and Lionel SPINELLI for usefull statistical advices. We thank for the help of Marc BARAD and Sylvain BIGOT from the CIML Cytometry core facility and Alice CARRIER and Laurence BORGE from the CRCM metabolomics core facility. I thank Pierre GOLSTEIN for mentoring and correcting the manuscript. The authors declare to have no competing interest.

## Authors’ contribution

Conceptualization, R.J.A.

Method development, R.J.A., A.C.

Validation, R.J.A., A.C, R.C, E.B., P.C., V.C., J.P.G.

Formal Analysis, R.J.A., A.C, B.S.

Investigation, R.J.A., A.C, M.F.K. and P.P.

Tissue Resources, E.T., D.F., S.B., M.F.K and P.P.

Writing – Original Draft, R.J.A.

Writing – R.J.A., A.C, E.G., M.F.K., and P.P.

Review & Editing, R.J.A., A.C, E.G, R.C., M.F.K. and P.P.

Visualization, R.J.A., A.C, E.G, M.F.K and P.P.

Supervision, R.J.A., M.F.K and P.P.

Funding Acquisition, R.J.A., M.F.K and P.P.

---

**Figure S1, Related to Figure 1.**
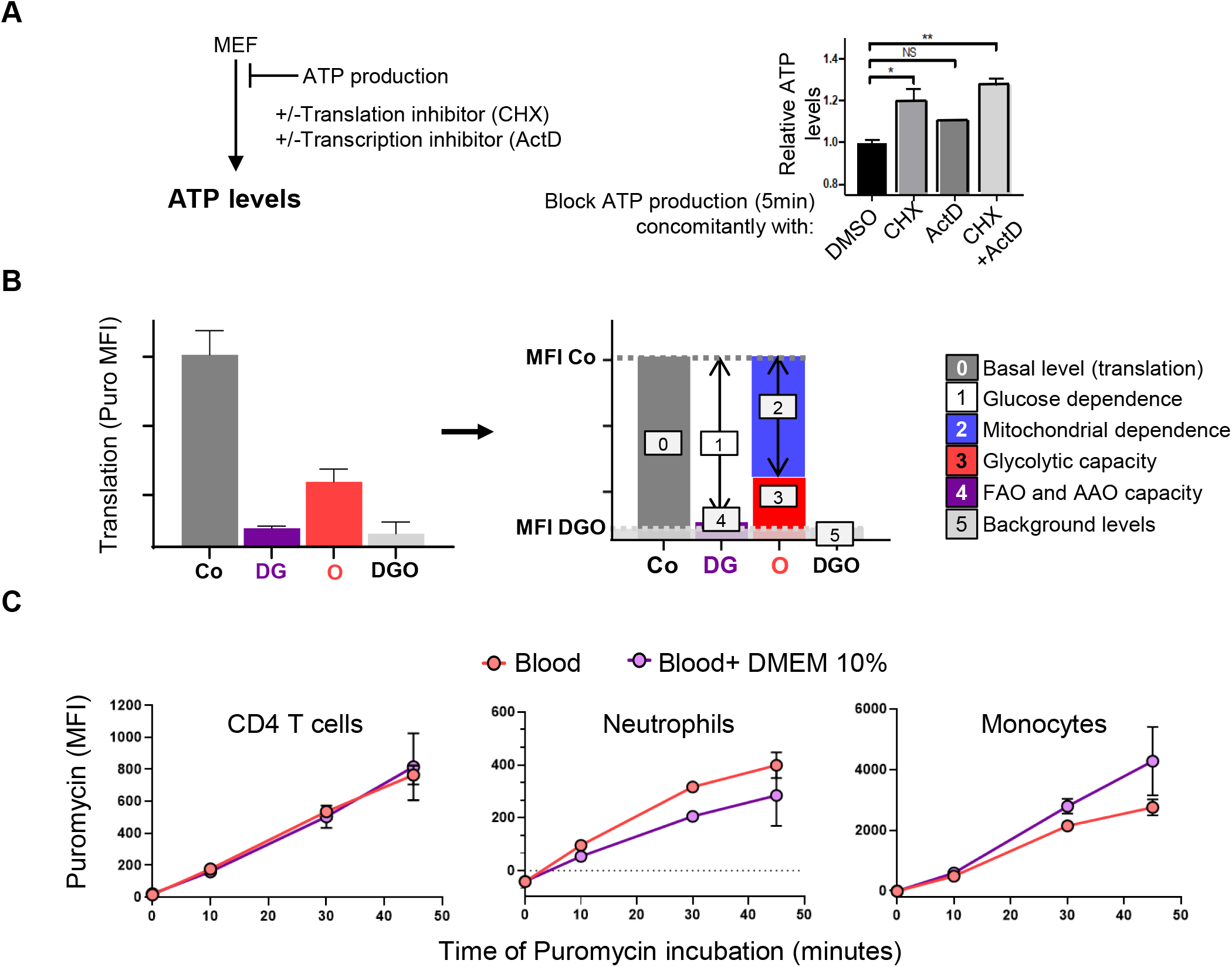
Translation, transcription and time of blood SCENITH. **A)** Impact of simultaneous block of ATP synthesis, translation (CHX), transcription (Actinomycin D, ActD) or both translation and transcription in the levels of ATP in MEFs (* p<0.05, ** p<0.01, N=3). **(B)** Representative scheme of SCENITH metabolic profile (left) and five metabolic parameters/measurements that can be extracted from the graph (right) using DG and O as inhibitors. **(C)** Level of puro staining from whole blood or whole blood mixed 1:4 with DMEM 10% FCS and incubated with puro for different amounts of time. Right panel, shows the translation levels (Y-axis) as a function of time of incubation with puro measured in different immune cells (N=3, each in duplicates).

**Figure S2, Related to Figure 2.**
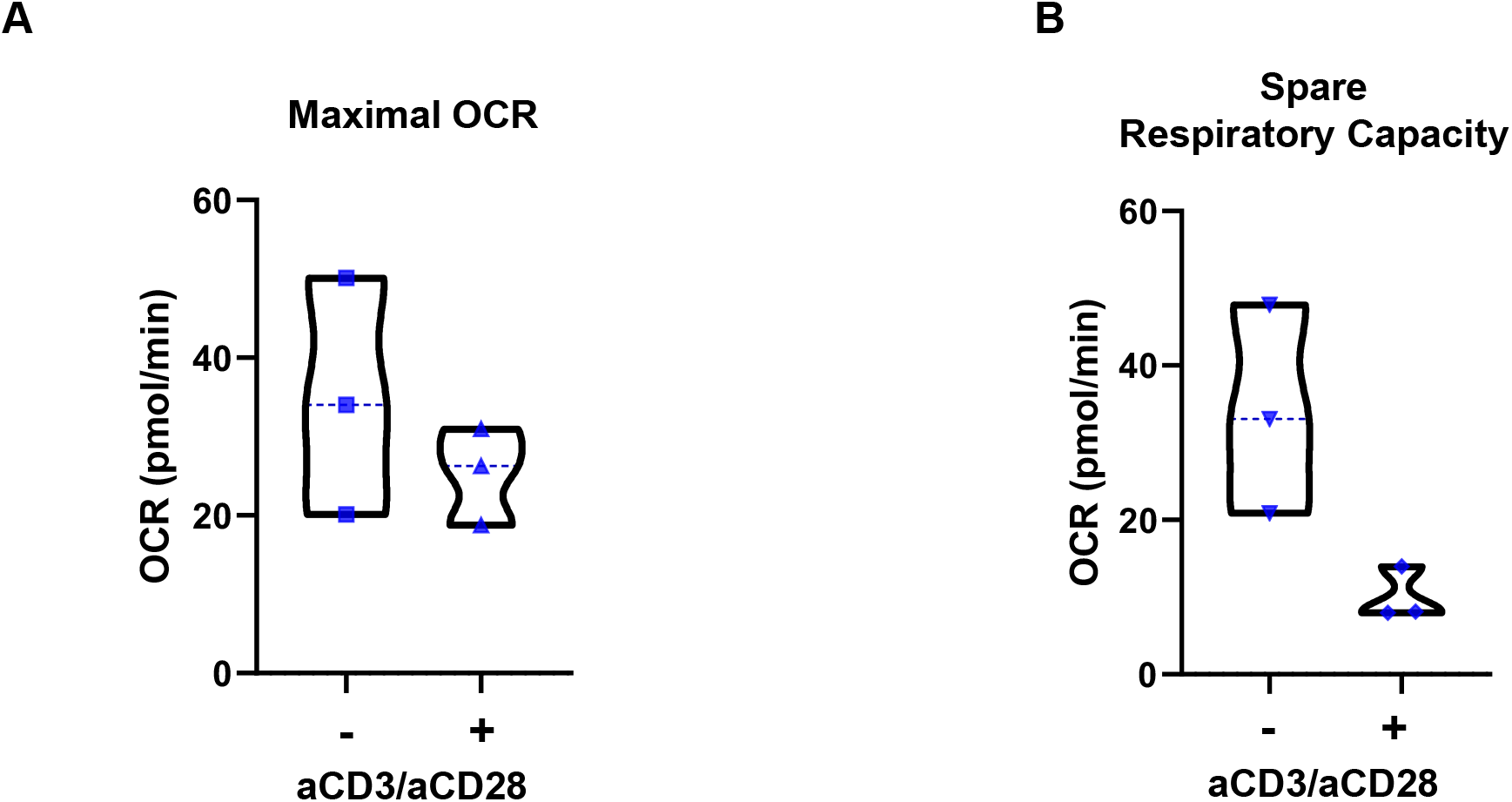
Seahorse metabolic parameters in non-activated and activated purified human T cells and SCENITH in whole blood. **(A)** Seahorse maximal oxygen consumption rate (OCR, upon FCCP) in non-activated or T cells stimulated over night with CD3/CD28 activator beads (N=3). **(B)** Seahorse spare respiratory capacity in non-activated or T cells stimulated over night with CD3/CD28 activator beads (N=3).

**Figure S3, Related to Figure 3.**
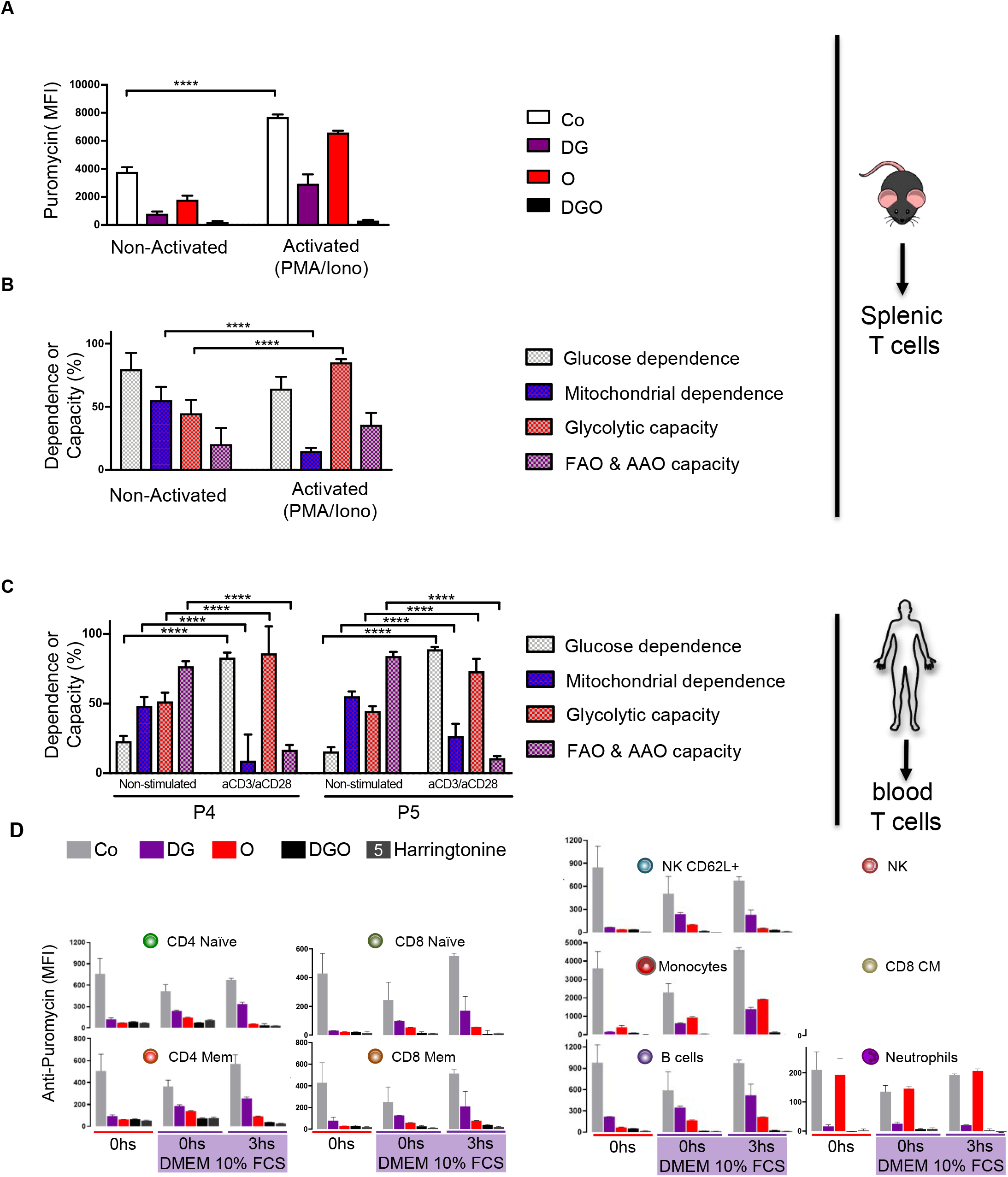
Changes in SCENITH EM profile in mouse and human T cells upon activation. **(A)** Mouse splenic T cells analyzed using SCENITH. Anti-Puro gMFI is indicated in CD3^+^CD8^+^CD4^−^ non-activated, left or PMA/Ionomycin-treated right (N=3, *** p<0.001) **(B)** EM profiles calculated from the values shown in (A). **(C)** EM profile of non-stimulated or stimulated human blood central memory CD4 T cells (CD3^+^CD4^+^CD45RO^+^CCR7^+^) from two subjects (P4 and P5) using the SCENITH method. N=3, ANOVA on at least 2 independent experiments (* p<0.05; ** p<0.005; ***p<0.0005). **(D)** Effect of short-term incubation of whole blood with cell culture media in the translation levels in immune cells. Pooled whole blood from three mice was pre-incubated with DMEM 10% FCS (0, or 3hs) or directly incubated with Co, DG, O, DGO or Harringtonine (translation initiation inhibitor) and Puro. Anti-Puro gMFI (experimental duplicates) is shown in the cell types comparing the response when treatments were applied directly on whole blood (0hs, left), on whole blood diluted in cell culture media (0hs, DMEM 10% FCS) or pre-incubated during 3hs with cell culture media (3hs DMEM 10 % FCS). Results shown are representative from one of the three independent experiments performed.

**Figure S4, Related to Figure 4.**
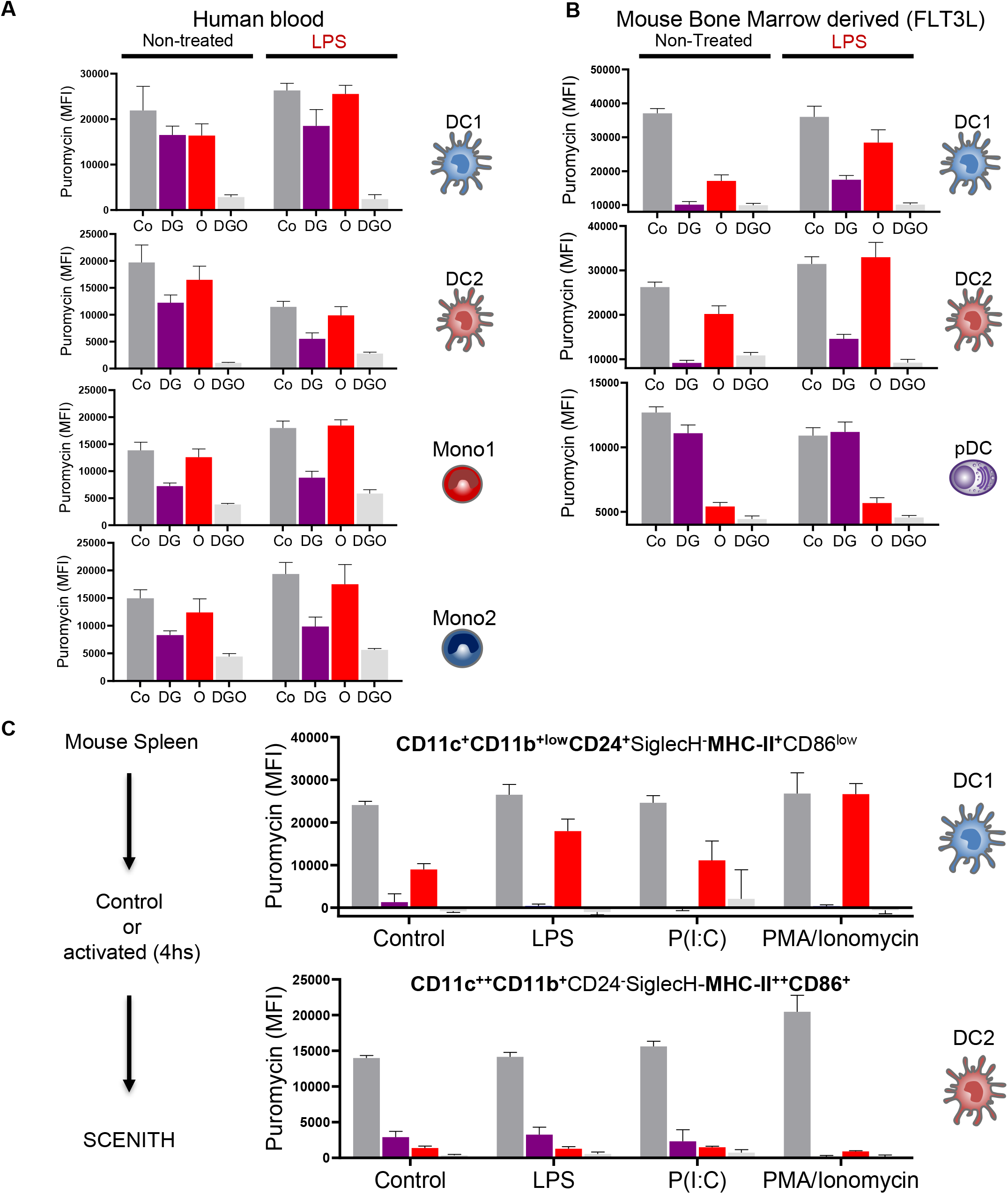
Metabolic profile of human and mouse myeloid cells. **(A and B)** SCENITH metabolic profile of human blood mononuclear cells (A) and mouse *in vitro* bone marrow derived (FLT3L-DC) (B) stimulated 4 hours or not with LPS. For both, DC1 are: CD11c^+^MHC-II^+^SIRPa^−^CD24^+^SiglecH^−^, DC2: CD11c^+^MHC-II^+^SIRPa^+^XCR1-CD24-SiglecH-, pDC: CD11c^+^MHC-II^+^SIRPa^−^CD24^−^SiglecH^+^ Mono1:Lin-CD14^+^CD16^−^MHC-II^+/-^ and Mono2: Lin-CD14^+^CD16^+^MHC-II^+^ (N=3, each in triplicates, two-way ANOVA, * p<0.05; ** p<0.005; ***p<0.0005). **(C)** Mouse splenocytes were treated for 4 hours with or without TLR ligands (LPS, 100ng/ml; Poly (I:C) 10ug/ml) or PMA/ionomycin (10 ng/ml PMA; 2.5 μM Ionomycin). SCENITH was performed and the resulting profile of protein synthesis levels in DC1 (top) or DC2 (bottom) are shown (N=3 independent spleens) (* p<0.05; ** p<0.005; ****p<0.0001).

**Figure S5, Related to Figure 5.**
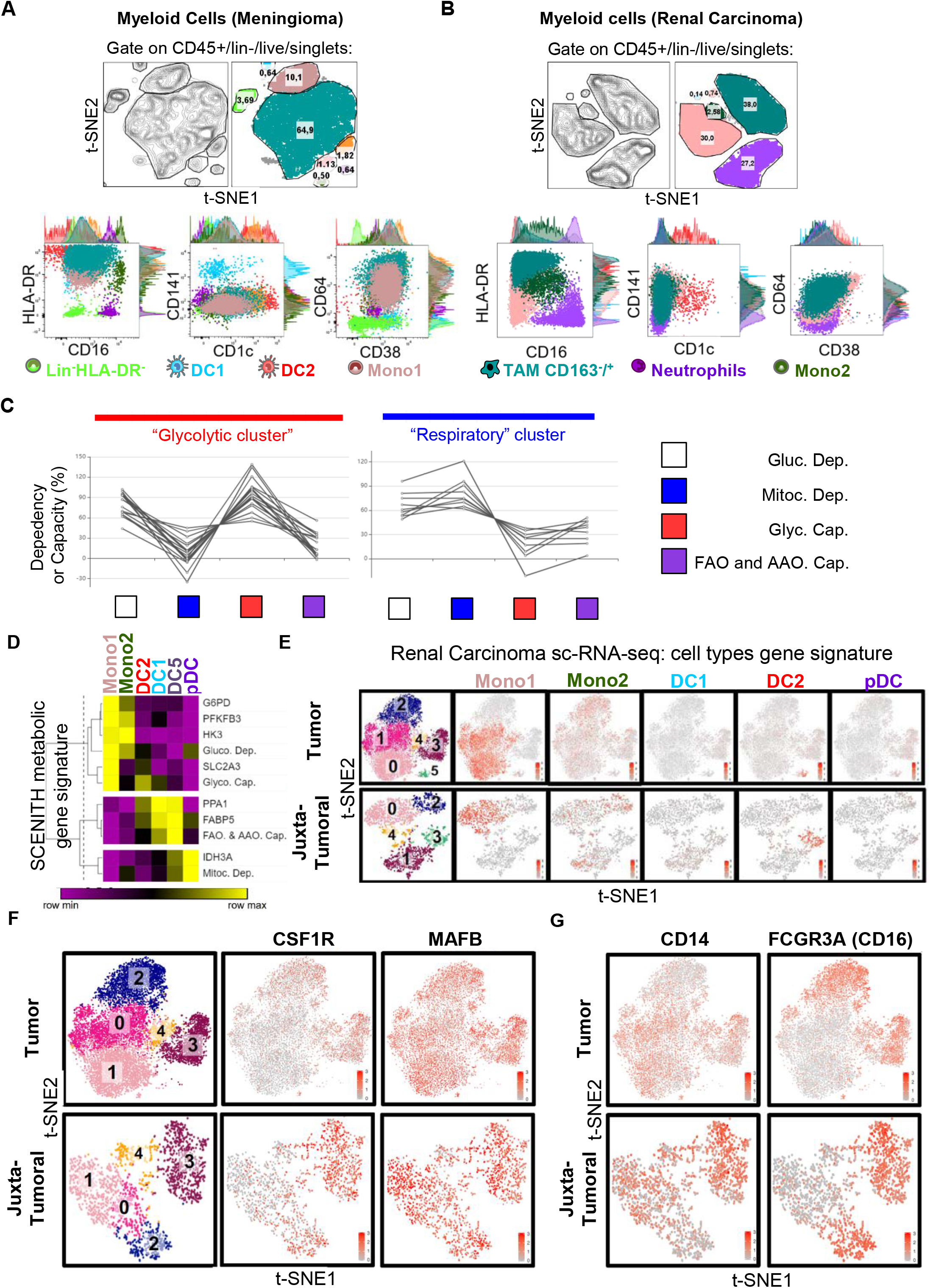
Composition, phenotype and metabolic profile of multiple immune cells populations can be analyzed in parallel in human tumors, *ex vivo* both by SCENITH and scRNAseq. **A and B)** t-SNE graph showing the myeloid cells analyzed by SCENITH in the meningioma (A), and renal carcinoma (B). Number in t-SNE means the percentage parent for each subpopulation and dot plot under t-SNE means the phenotype in each subpopulation (N=4). **(C)** Clustering and identification of two distinct metabolic clusters of cells. The metabolic profile of each subset from each tissue was used to cluster cells. Two clusters were observed with different metabolic profile, and were name accordingly as Glycolytic cluster and Respiratory cluster of cells (N=4). **(D)** Heatmap of metabolic gene expression and SCENITH functional metabolic profile in myeloid cells from human PBMC (N=5). **(E-G)** Far left, t-SNE display and graph-based clustering of CD3^−^CD19^−^CD20^−^CD56^−^, HLA-DR^+/-^ myeloid cells sorted from Renal carcinoma (top) or Juxta tumoral tissue (bottom) biopsies and processed for scRNAseq. From left to right, level of expression of (grey=low, red=high) gene signatures of blood myeloid cell types **(E)**(Villani et al. 2017); level of CSF1R and MAFB **(F)** and CD14 and CD16 **(G)** in myeloid cells clusters observed in tumor (upper line) and juxta-tumoral tissue.

**Figure S6, Related to Figure 5.**
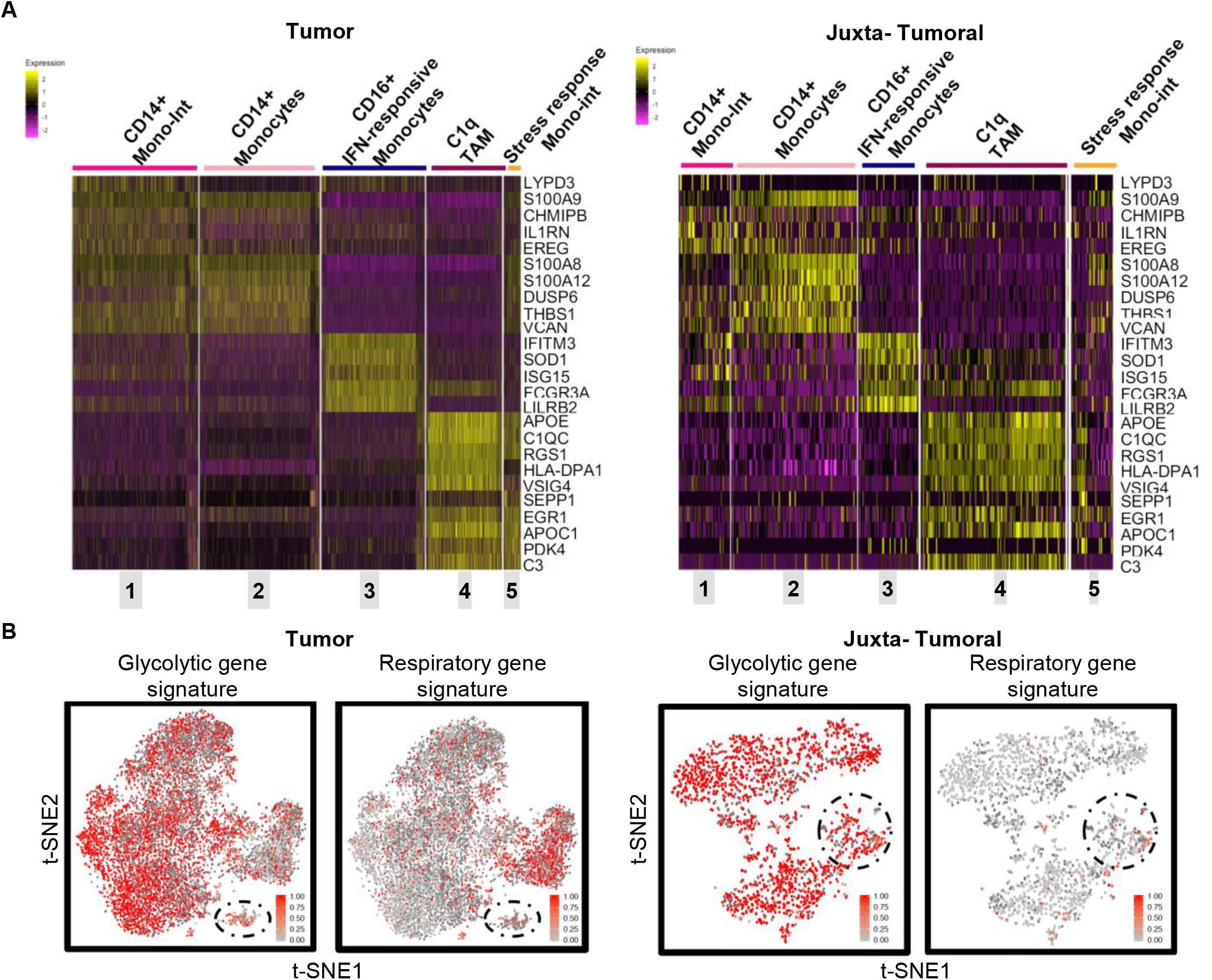
**(A)** Heatmap displaying top 5 differentially expressed genes for each cluster of monocytes and macrophages present in renal carcinoma (left) or in juxta-tumoral tissue (right) when comparing cluster 0 through 4 in renal carcinoma (ranked by log fold change). **(B)** SCENITH derived Glycolytic (left) and Respiratory (right) gene signature expression distributed across t-SNE plot of myeloid cells sorted from renal carcinoma (top) or juxta-tumoral tissue (bottom) with DC clusters circled.

**Table S1, Related to Table 1.** Comparative table of methods to profile metabolism.

| Method | CyTOF (e.g. Met-Flow, scMEP) | MSI | Seahorse® | SCENITH™ |
| --- | --- | --- | --- | --- |
| Output | Metabolic phenotype | Metabolomic profile (unbiased) | Metabolic capacities and dependencies | Metabolic capacities and dependencies |
| Functional profile of the cells (# of treatments) | NO | NO | YES (4) | YES (non-limited) |
| Cell purification required | NO | NO | YES | NO |
| Single cell resolution | YES | YES | NO | YES |
| Fixation compatible | Yes | YES | NO | YES |
| Stability of the readout | High | Variable | Low | High |
| Phenotypic analysis | YES | NO | NO | YES (including Metabolic phenotype) |
| Metabolite profiling | NO | YES | NO | NO |
| Compatible with cell sorting | NO | NO | NO | YES* |
| *In vivo* application | NO | NO | NO | NO |
| *Ex vivo* application | YES | YES | NO | YES |
| *In vitro* application | YES | YES | YES | YES |
| Metabolic Readout | Levels of markers (min 10 channels) | Metabolite levels | Changes in extracellular pH and [O<sub>2</sub>] | Changes in Protein synthesis levels (one channel). |
| Time (Hs) from sampling to metabolic profiling | NA | NA | 24 | 0-1 |
| Total time | 6 | 6 | 24 | 4-7 |
| # cells required in subsets | 500 | 200 | 1.000.000 | 2000 |
| Accessibility | Low | Low | Low | High |
| Equipment needed | CyTOF cytometer | Any Imaging Mass cytometer | Seahorse Analyzer | Any Flow cytometer<sup>#</sup> |

\* Not shown

**Table S2.** Related to STAR Methods SCENITH. Troubleshooting table.

| Problem | Potential cause | Suggestion |
| --- | --- | --- |
| **Signal for translation (Puro MFI) is too low or lower signal in control treatment** | Cells were not kept at 37°C during treatments. | Prepare all reagents and equilibrate the temperature. Neither put cells on ice, nor keep them at room temperature. |
|  | Puro stock was not correctly stored. | Avoid exposure to light and freeze thawing cycles. Use small aliquots of puro and discard after each use. |
|  | An inappropriate concentration of puro was used | Puro concentration can be cell dependent and should be determined experimentally for each sample being tested. too high, or too low concentrations result in low staining. |
|  | The incubation time with puro was insufficient | Do a kinetic experiment of puro incubation time ranging from 5-15-30-45 min. |
|  | Contaminated cells | Obtain fresh culture with fresh medium to prevent contamination. 1% Penicillin (not streptomycin) can be used |
|  | Puro resistance in your cells | Change antibiotic resistance into the construct to be compatible with SCENITH™. |
|  | Cell culture medium was depleted of nutrients. | Change media 2~8 hr before the experiment. |
|  | Too many cells during staining | Do not stain in each tube more than 5 million cells. Adapt the volume of antibody. |
| **Signal for translation (Puro MFI) is too high** | Commercially available clones can show substantial background. | Ask for the SCENITH™ kit containing the R4743L-E8 mAb clone. www.scenith.com |
| **Signal for translation (Puro MFI) is higher in my cells treated with inhibitor than the control** | Some metabolic inhibitors make cells to switch to other sources to compensate. The activation of this compensatory mechanism might result in higher levels of translation than the control. | If cells show increased translation upon treatment when calculating the metabolic dependency, the formula would result in a negative dependency value. Conceptually a negative dependency is equal to no dependency (0). |
|  |  | As the inhibitor induce an increase in the translation levels and cells are only relying on the compensatory pathway, then the formula will reveal a percentual metabolic capacity that is over 100%. Meaning that cells have the metabolic capacity to sustain even higher levels of translation than control cells. |
| **Signal of cell surface markers decrease after treatment with inhibitors** | An inappropriate time of treatment with inhibitors was performed | Time of treatment with inhibitors depends on the metabolic activity of the cells and have to be determined experimentally. Reduce the time of DGO treatment (i.e. 10min DG, then add O for 5min, then Puro). Use H treatment as negative control present in the kit. |
|  |  | Perform intracellular staining of internalized markers. |
| **Signal is present in DGO (all inhibitors)** | Oligomycin acts faster than DG (competitive inhibitor of glucose). | Treat cells 10min with 2DG and then add the Oligomycin for 5min before puro treatment, and use that measure as DGO treatment. |
|  | High background from cells | Use Fc block, Fcs 2% or BSA 1-3% during staining can reduce background |
| **Signal doesn't change with any kind of treatment** | Inhibitor doesn't work | Use inhibitors at 37°C to have powerful effect |
|  | The concentration and/or time of treatment are not optimized | Inhibitors concentration can be cell dependent and should be determined experimentally for each cell line being tested. |
| **Lot of doublet cells during FACS acquisition** | Cells were aggregated | Obtain a single cell suspension by using 2mM EDTA in FACS buffer |

## Notes

### Competing Interest Statement

The authors declare to have no competing interest. There are restrictions to the commercial use of SCENITH due to pending patent application (PCT/EP2020/060486).

